# Relief of allosteric inhibition, redox imbalance, and transport limitations enables high-yield L-malate production in *Escherichia coli*

**DOI:** 10.64898/2026.05.04.722580

**Authors:** Moses Onyeabor, Lizbeth M. Nieves, Gavin Kurgan, Junpei Xiao, Logan Kurgan, Brittany Retallack, Haiwei Gu, Xuan Wang

**Affiliations:** School of Life Science, Arizona State University, Tempe, AZ, USA; College of Health Solutions, Arizona State University, Phoenix, AZ 85004, USA

## Abstract

Malic acid is a C4 dicarboxylic acid traditionally produced from petroleum and widely used in the food industry. As a sustainable alternative, it can also be produced as a value-added platform chemical from biomass. Previously, the *Escherichia coli* strain XZ658 was engineered to produce L-malate via the carbon-fixation reductive branch of the TCA cycle. In this study, we further improved this system by relieving allosteric regulation of citrate synthase, addressing redox imbalance, and enhancing malate export. These modifications approximately doubled the L-malate titer in the final strain MO128 compared to XZ658 under simple batch fermentation conditions. The process achieved a high mass yield of 1.2 g malate g⁻¹ glucose, highlighting the carbon-fixation capacity of the reductive TCA pathway for fermentative malate production.

## 1. Introduction

Microbial conversion of renewable feedstocks, such as lignocellulosic biomass, into fuels and chemicals through fermentation-based manufacturing processes has been widely studied in the past decades as viable ecofriendly alternative to fossil-based production (Nieves et al., 2015). The top 12 value-added chemicals derived from biomass with the potential to replace petrochemicals were identified by the Department of Energy (Werpy, 2004). 1, 4-C4 dicarboxylates such as succinate, fumarate, L-malate, and L-aspartate are among these identified top commodity chemicals with a multibillion-dollar market value (Wei et al., 2021; Wu et al., 2025). Currently, the major routes to produce these compounds are still based on the petrochemical platform (Yin et al., 2015). Although there are many examples in scientific literature regarding metabolic engineering of microbes to produce valuable renewable chemicals including 1, 4-C4 dicarboxylates, the low production performance (titer, yield and productivity) limits the economical large-scale production of these chemicals (Wei et al., 2021; Wu et al., 2025).

1, 4-C4 dicarboxylates such as succinate and L-malate are central metabolites without significant cellular toxicity even at high concentrations (Wu et al., 2025). Microbial production of these compounds has a high theoretical yield due to the carbon-fixation step of the TCA reductive branch during their biosynthesis. For example, the maximal theoretical yields for succinate and malate are 1.12 and 1.49 g per g glucose, respectively (Lin et al., 2005; Zhang et al., 2011). A wide range of microorganisms including both fungi and bacteria were developed to produce malate using aerobic or microaerobic processes (Khandelwal et al., 2023; Wei et al., 2021; Xu et al., 2024). Although high titers (100 g liter^-1^ or higher) have been achieved in fungal hosts such as *Aspergillus. oryzae* (Brown et al., 2013), *Aureobasidium pullulans* (Zou et al., 2013), *Pichia kudriavzevii* (Xi et al., 2023), and *Myceliophthora thermophila* (Zhang et al., 2011, 2025), the yields often remain low and the complica gtion (Xu et al., 2024). In addition, filamentous fungi such as *A. oryzae* and *M. thermophila* present challenges for large-scale fermentation processes. The use of CaCO₃ in fungal malic acid production also creates downstream processing issues due to gypsum waste generation. Using non-filamentous, unicellular microbes, along with soluble bases and inorganic carbon sources such as potassium bicarbonate as co-substrates, may enable more straightforward process development for malic acid production.

In previous work, we successfully engineered *E. coli* for malate production using a precursor strain with enhanced PCK activity and multiple targeted gene deletions to direct metabolic flux toward malate (Zhang et al., 2011). Homo-malate fermentation is theoretically a redox-balanced process with net ATP generation when phosphoenolpyruvate (PEP) carboxykinase (Pck) is used instead of the native PEP carboxylase (Ppc) (Zhang et al., 2011, 2009). Production of each malate molecule requires the fixation of one CO₂/HCO₃⁻ while preserving all carbon from glucose, allowing a theoretical yield of up to 2 moles of malate per mole of glucose, corresponding to a mass yield of 1.49 g malate g⁻¹ glucose (Fig.1A). However, the slow growth, low yield of cell mass and low production metrics suggest that intrinsic metabolic constraints limit the carbon-fixation capacity of the malate production under oxygen-limiting fermentative conditions. The carbon fixation capacity of the TCA reductive branch is not fully exploited in this strain.

**Figure 1.**
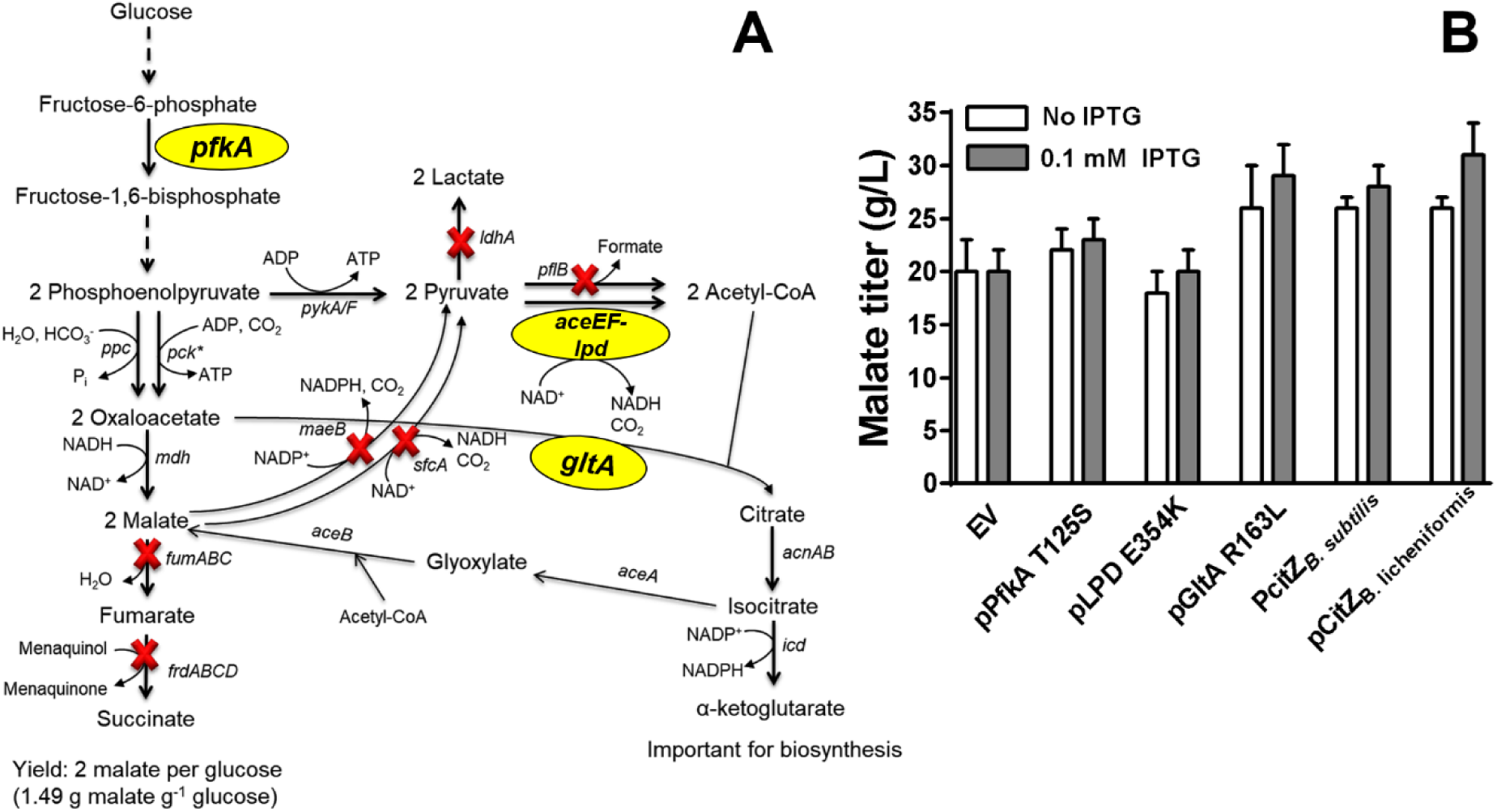
The metabolic pathway for L-malate fermentative production and the investigation of potential allosteric regulatory sites. A) Potential metabolic allosteric regulation for malate production. The reductive branch of the TCA cycle was engineered to accumulate malate as the sole fermentation product by deleting genes encoding fumarases (*fumB* and *fumAC*), fumarate reductase (*frdABCD*), and malic enzymes (*maeB* and *sfcA*). A redox-balanced malate-producing pathway converts one molecule of glucose into two molecules of malate. Some intermediate steps in glycolysis are not shown for clarity and simplicity. The malate-producing strain contains spontaneous mutations in *pck* and *ptsI* that were important to enhance carbon flow in the reductive branch of TCA cycle (Zhang et al., 2011). Three potential allosteric regulation sites are indicated in ovals: 6-phosphofructokinase, pyruvate dehydrogenase, and citrate synthase encoded by *pfkA*, *aceEF-lpd* and *gltA*, respectively. ‘X’ in bold indicates the chromosomal inactivation of the designated genes. Abbreviation: acetyl-CoA, acetyl-Coenzyme A. B) Effect of different plasmids on malate production in XZ658. Fermentations (n ≥3) with and without IPTG were carried out in NBS mineral salts medium with 50 g liter^-1^ glucose and 100 mM potassium bicarbonate for 144 hours (37 °C, pH 7.0). *p < 0.05 as estimated by one-tailed Student’s t test relative to corresponding empty vector (EV) controls.

Allosteric inhibition often limits microbial production from achieving desired performance metrics. For allosteric regulation associated with malate production, three major regulatory sites are involved in C4 dicarboxylate biosynthesis pathways: 6-phosphofructokinase (Pfk) (Babul et al., 1977), lipoamide dehydrogenase (Lpd) (Kim et al., 2008), and citrate synthase (GltA) (Danson et al., 1979). Pfk catalyzes the irreversible phosphorylation of fructose-6-phosphate in an early step of glycolysis. Lpd is the E2 component of the pyruvate dehydrogenase complex, which catalyzes the decarboxylation of pyruvate to form acetyl-CoA. GltA catalyzes the condensation of oxaloacetate and acetyl-CoA to form citrate in the TCA cycle, representing the first committed step toward α-ketoglutarate production, an important precursor for amino acid biosynthesis (Underwood et al., 2002). In *E. coli*, PEP is a negative allosteric regulator for Pfk and high intracellular levels of NADH strongly inhibits both Lpd and GltA (Babul et al., 1977; Danson et al., 1979; Kim et al., 2008). This allosteric inhibition can be mitigated by replacing native enzymes with heterologous isozymes or engineered mutants that are resistant to inhibition (Auzat et al., 1994; Kim et al., 2008; Stokell et al., 2003; Underwood et al., 2002).

In this study, we identified allosteric inhibition of citrate synthase, high NADH/NAD⁺ ratio, and product export as significant metabolic constraints for malate production in XZ658. Relieving these metabolic constraints increased malate titer and yield by 105% and 61%, respectively, under simple batch fermentation conditions. The high yield achieved for microbial malate production highlights the carbon-fixation potential of the reductive branch of the TCA cycle under anaerobic fermentation.

## 2. Materials and methods

### 2.1. Strains and plasmids

Strains and plasmids used in this study are listed in Table 1. XZ658 was kindly provided by Dr. Lonnie Ingram’s group at University of Florida (Zhang et al., 2011). Other strains were constructed using λ-red recombinase-mediated two-step method as previously described (Sievert et al., 2017). To express heterologous or mutant genes resistant to allosteric regulation from a plasmid, the native ribosomal binding sites, the open reading frames, and terminator regions were cloned into pTrc99A under the control of the isopropyl β-D-1-thiogalactopyranoside (IPTG) inducible *trc* promoter. The construction details such as used restriction sites are described in Table 1.

**Table 1.**
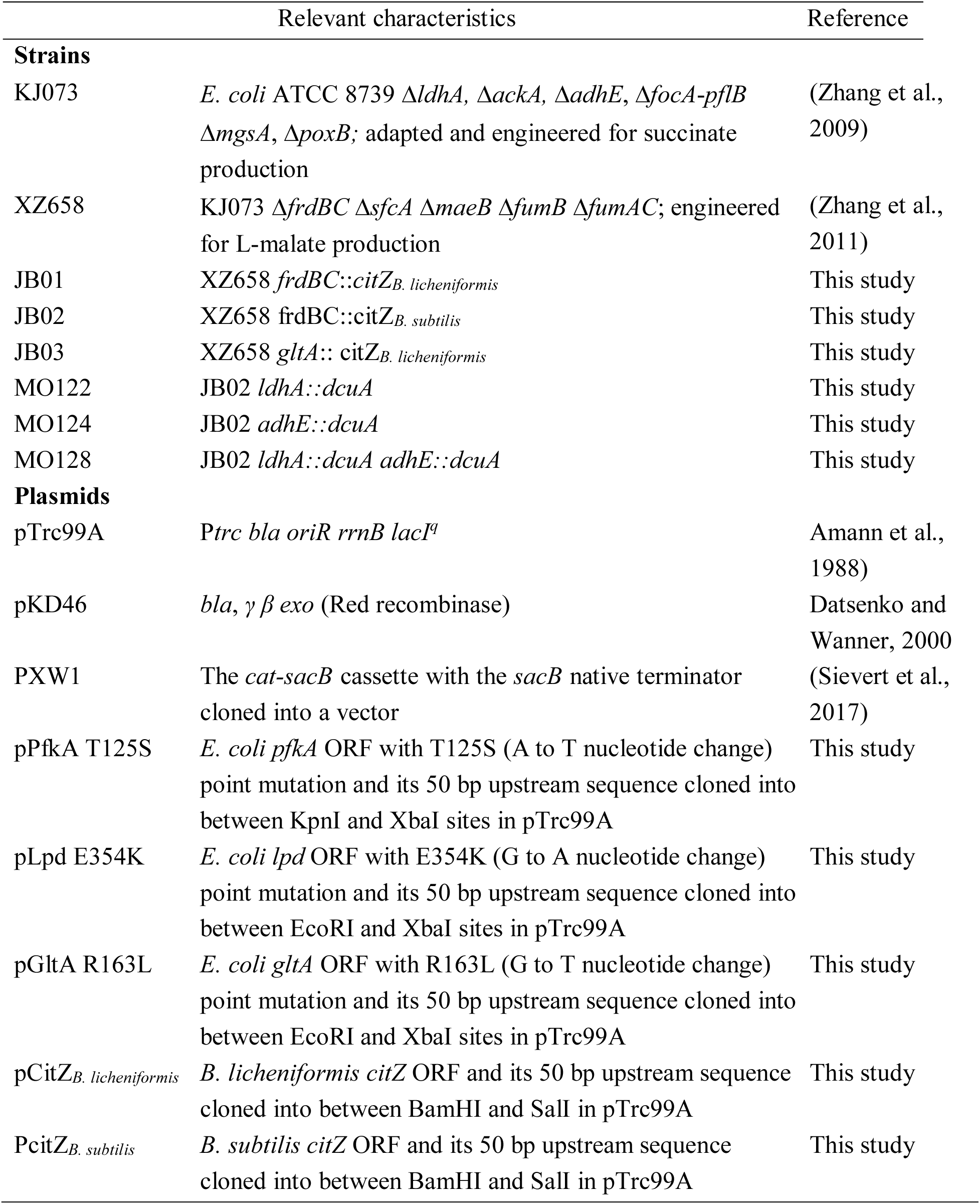
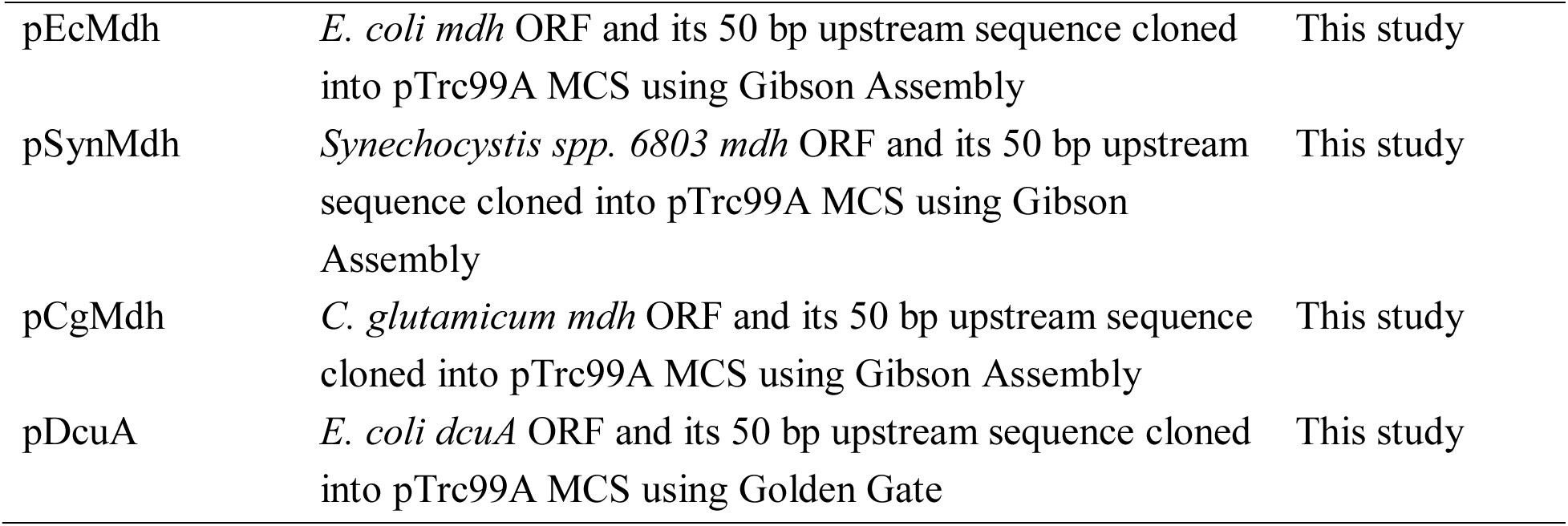
Strains and plasmids used in this study.

|  | Relevant characteristics | Reference |
| --- | --- | --- |
| **Strains** |  |  |
| KJ073 | *E. coli* ATCC 8739 $\Delta ldhA$, $\Delta ackA$, $\Delta adhE$, $\Delta focA$-*pflB* $\Delta mgsA$, $\Delta poxB$; adapted and engineered for succinate production | (Zhang et al., 2009) |
| XZ658 | KJ073 $\Delta frdBC$ $\Delta sfca$ $\Delta maeB$ $\Delta fumB$ $\Delta fumAC$; engineered for L-malate production | (Zhang et al., 2011) |
| JB01 | XZ658 *frdBC::citZ<sub>B. licheniformis</sub>* | This study |
| JB02 | XZ658 *frdBC::citZ<sub>B. subtilis</sub>* | This study |
| JB03 | XZ658 *gltA::citZ<sub>B. licheniformis</sub>* | This study |
| MO122 | JB02 *ldhA::dcuA* | This study |
| MO124 | JB02 *adhE::dcuA* | This study |
| MO128 | JB02 *ldhA::dcuA adhE::dcuA* | This study |
| **Plasmids** |  |  |
| pTrc99A | *P<sub>trc</sub> bla oriR rrnB lacI<sup>q</sup>* | Amann et al., 1988) |
| pKD46 | *bla*, $\gamma$ $\beta$ *exo* (Red recombinase) | Datsenko and Wanner, 2000 |
| PXW1 | The *cat-sacB* cassette with the *sacB* native terminator cloned into a vector | (Sievert et al., 2017) |
| pPfkA T125S | *E. coli* *pfkA* ORF with T125S (A to T nucleotide change) point mutation and its 50 bp upstream sequence cloned into between KpnI and XbaI sites in pTrc99A | This study |
| pLpd E354K | *E. coli* *lpd* ORF with E354K (G to A nucleotide change) point mutation and its 50 bp upstream sequence cloned into between EcoRI and XbaI sites in pTrc99A | This study |
| pGltA R163L | *E. coli* *gltA* ORF with R163L (G to T nucleotide change) point mutation and its 50 bp upstream sequence cloned into between EcoRI and XbaI sites in pTrc99A | This study |
| pCitZ<sub>B. licheniformis</sub> | *B. licheniformis* *citZ* ORF and its 50 bp upstream sequence cloned into between BamHI and SalI in pTrc99A | This study |
| PcitZ<sub>B. subtilis</sub> | *B. subtilis* *citZ* ORF and its 50 bp upstream sequence cloned into between BamHI and SalI in pTrc99A | This study |
| pEcMdh | *E. coli mdh* ORF and its 50 bp upstream sequence cloned into pTrc99A MCS using Gibson Assembly | This study |
| pSynMdh | *Synechocystis spp. 6803 mdh* ORF and its 50 bp upstream sequence cloned into pTrc99A MCS using Gibson Assembly | This study |
| pCgMdh | *C. glutamicum mdh* ORF and its 50 bp upstream sequence cloned into pTrc99A MCS using Gibson Assembly | This study |
| pDcuA | *E. coli dcuA* ORF and its 50 bp upstream sequence cloned into pTrc99A MCS using Golden Gate | This study |

### 2.2. Media and growth conditions

During strain construction, cultures were grown aerobically at 30°C, 37°C, or 39°C in Luria-Bertani broth as needed. NBS (New Brunswick Scientific) mineral salts medium supplemented with 50 g liter^-1^ glucose, 10 mM acetate and 100 mM potassium bicarbonate was used in batch fermentation as previously described (Zhang et al., 2011). Ampicillin (50 mg liter^-1^), kanamycin (50 mg liter^-1^), or chloramphenicol (40 mg liter^-1^) were added as needed.

### 2.3. Genetic methods

Red recombinase technology was used to perform seamless chromosomal deletion and gene replacement as previously described (Sievert et al., 2017). The DNA fragments used for the first-step integration were obtained by PCR to amplify the *cat-sacB* cassette from plasmid pXW1 and primers provided 50 bp or longer homologous sequences to facilitate homologous recombination. The DNA fragments used for the second-step integration containing homologous regions (∼500 bp flanking sequence on each side) were assembled by fusion PCR (Higuchi et al., 1988). Point mutations in target genes were also introduced using fusion PCR with the relevant primers listed in Table S1. During strain construction, 5% (w/v) arabinose and 10% (w/v) sucrose were used for induction of λ-Red recombinase synthesis and for *sacB* counterselection, respectively. Positive clones from genetic manipulation were verified by colony PCR and Sanger sequencing of the PCR-amplified target chromosomal regions.

### 2.4. Fermentation and analysis

All batch fermentations were performed using NBS (New Brunswick Scientific) mineral salts medium supplemented with 50 g liter^-1^ glucose, 100 mM potassium bicarbonate and 10 mM sodium acetate in small fermentation vessels with 300 ml as the working volume as previously described (Zhang et al., 2011). Pre-inoculum for fermentations was started by transferring fresh colonies from NBS 2% glucose (w/v) plates into a shake flask (100 ml NBS medium, 2% glucose). After 16 h (37°C, 120 rpm), this culture was diluted into a fermentation vessel containing 300 ml NBS medium to provide an inoculum of 0.044 g cell dry wt (CDW) liter^-1^ (an optical density value at 550nm of 0.1) and batch fermentations were performed for 144 h. Batch fermentations of strains with plasmids were operated in a similar manner with the supplementation of 50 mg liter^-1^ ampicillin and IPTG at the appropriate concentrations to induce the expression of tested genes. All fermentations were maintained at pH 7.0 by the automatic addition of base solution (2.4 M potassium carbonate and 1.2 M potassium hydroxide) as previously described (Zhang et al., 2011).

Glucose and organic acids in fermentation broth were measured by high-performance liquid chromatography (HPLC) using an Aminex® HPX-87H column (Bio-Rad) and 4 mM sulfuric acid as the mobile phase (Sievert et al., 2017; Zhang et al., 2011). Cell dry weight was derived from the measured optical density at 550 nm. Experimental data represent an average of three or more measurements with standard deviations.

### 2.5. Aqueous metabolite extraction and measurement

Cells from fermentation vessels were harvested at the exponential growth phase for metabolite extraction and quantitation using gas chromatography–mass spectrometry (GC-MS) analysis as previously described (Kurgan et al., 2019). Measurement of intracellular metabolites was performed by the Arizona Metabolomics Laboratory at Arizona State University. GC-MS was performed with a general procedure from the Agilent Fiehn GC-MS Metabolomics RTL library as previously described (Kurgan et al., 2019). Metabolite levels were normalized to protein content, and the protein concentrations of cell lysates were measured using the BCA protein assay kit (Thermo Scientific, Rockford, IL). Intracellular concentrations of the pyridine nucleotides NADH and NAD⁺ were determined using the EnzyChrom™ NAD⁺/NADH assay kit (BioAssay Systems, Hayward, USA) according to the manufacturer’s instructions.

## 3. Results

### 3.1. Addressing allosteric inhibition in a malate-producing *E. coli* strain

The genes encoding *pfkA*, *lpd* and *gltA* variants resistant to allosteric inhibition were cloned into pTrc99A with their native ribosomal binding sites and terminator regions (Table 1). The missense mutations of T125S in *pfkA* and E354K in *lpd* were used to release the allosteric regulation of 6-phosphofructokinase and pyruvate dehydrogenase in a previously engineered malate producer XZ658 (Zhang et al., 2011). There are two types of bacterial citrate synthases with distinct subunit arrangements and allosteric regulatory properties: the type I enzymes in Gram-positive bacteria and archaea, and type II enzymes in Gram-negative bacteria (Weitzman, 1967). NADH strongly inhibits the type II citrate synthase in *E. coli* but has no effect on type I enzymes (Stokell et al., 2003; Weitzman, 1967). To release allosteric inhibition of citrate synthase, two different kinds of variants insensitive to NADH inhibition were used in this study: i) the mutant of *E. coli* citrate synthase (GltA R163L)(Stokell et al., 2003) and ii) the citrate synthase CitZ from *Bacillus subtilis* and *Bacillus licheniformis* (Underwood et al., 2002).

Overexpression of *pfkA* and *lpd* mutants upon IPTG induction had essentially no impact on malate production compared to empty vector (EV; pTrc99A) control (Fig. 1B and Table 2). In contrast, induced expression of either the *gltA* mutant or *citZ* genes from both *Bacillus* strains increased malate titer and yield ranging from 40 to 55% and 45 to 60%, respectively. Even without induction, the leaky expression of these variant genes led to increased malate titers and yields (Fig. 1B and Table 2). This improvement of malate production is correlated to the increased biomass yield and sugar consumption (Table 2). Titers of side products, especially the main side product D-lactate, are similar to the EV control and thus more sugar conversion into malate led to an increase in malate yield (∼ 1.0 g malate g^-1^ glucose used) (Table 2). This result suggests that allosteric inhibition of citrate synthase limits cell growth and malate production in XZ658.

**Table 2.** Effects of different plasmids on malate production in XZ658.

| Plasmids | Maximum Cell mass (g liter<sup>-1</sup>) | Glucose used (g) | Malate yield (g g<sup>-1</sup>)<sup>c</sup> | Productivity (g liter<sup>-1</sup> h<sup>-1</sup>) | Fermentation products (g liter<sup>-1</sup>)<sup>d</sup> |  |  |  |  |
| --- | --- | --- | --- | --- | --- | --- | --- | --- | --- |
|  |  |  |  |  | Mal | Lac | Pyr | Ace | For |
| pTrc99A<sup>a</sup> | 0.80 ± 0.10 | 28 ± 4 | 0.72 | 0.14 | 20 ± 3 | 8.4 ± 1 | 3.3 ± 0.7 | 0.86 ± 0.2 | 1.5 ± 0.1 |
| pTrc99A<sup>b</sup> | 0.75 ± 0.09 | 29 ± 1 | 0.69 | 0.14 | 20 ± 2 | 8.5 ± 0.6 | 2.0 ± 0.1 | 0.51 ± 0.2 | 1.4 ± 0.3 |
| pPfkA T125S<sup>a</sup> | 0.84 ± 0.09 | 29 ± 2 | 0.76 | 0.15 | 22 ± 2 | 10 ± 2 | 2.2 ± 0.7 | 0.99 ± 0.1 | 1.5 ± 0.1 |
| pPfkA T125S<sup>b</sup> | 0.70 ± 0.2 | 31 ± 4 | 0.74 | 0.16 | 23 ± 2 | 9 ± 2 | 2.0 ± 0.1 | 0.81 ± 0.1 | 1.2 ± 0.1 |
| pLpd E354K<sup>a</sup> | 0.70 ± 0.2 | 24 ± 1 | 0.75 | 0.12 | 18 ± 2 | 8.2 ± 0.7 | 1.8 ± 0.4 | 0.92 ± 0.2 | 1.4 ± 0.2 |
| pLpd E354K<sup>b</sup> | 0.80 ± 0.2 | 25 ± 3 | 0.8 | 0.14 | 20 ± 2 | 7 ± 2 | 2.2 ± 0.1 | 1.1 ± 0.6 | 1.4 ± 0.7 |
| pGltA R163L<sup>a</sup> | 1.1 ± 0.1 | 35 ± 3 | 0.74 | 0.18 | 26 ± 4 | 10 ± 3 | 3.6 ± 0.6 | 1.2 ± 0.4 | 2.3 ± 0.2 |
| pGltA R163L<sup>b</sup> | 0.90 ± 0.1 | 29 ± 2 | 1.0 | 0.2 | 29 ± 3 | 7.2 ± 0.4 | 3.0 ± 1 | 1.0 ± 0.4 | 2.1 ± 0.3 |
| pCitZ<sub>B. subtilis</sub><sup>a</sup> | 0.70 ± 0.09 | 25 ± 2 | 1.0 | 0.18 | 26 ± 1 | 6.1 ± 0.9 | 2.5 ± 0.1 | 0.40 ± 0.1 | 1.4 ± 0.1 |
| pCitZ<sub>B. subtilis</sub><sup>b</sup> | 0.92 ± 0.09 | 27 ± 4 | 1.0 | 0.19 | 28 ± 2 | 8.0 ± 1 | 2.9 ± 0.6 | 0.50 ± 0.1 | 1.4 ± 0.1 |
| PcitZ<sub>B. licheniformis</sub><sup>a</sup> | 0.97 ± 0.09 | 26 ± 5 | 1.0 | 0.18 | 26 ± 1 | 8.6 ± 0.2 | 3.4 ± 0.8 | 0.60 ± 0.1 | 1.6 ± 0.1 |
| PcitZ<sub>B. licheniformis</sub><sup>b</sup> | 0.90 ± 0.09 | 28 ± 3 | 1.1 | 0.22 | 31 ± 3 | 8.7 ± 0.5 | 3.7 ± 0.4 | 0.60 ± 0.1 | 1.4 ± 0.1 |

<sup>a</sup> Fermentations (n ≥ 3) without IPTG as controls were carried out in NBS mineral salts medium with 50 g liter<sup>-1</sup> glucose and 100 mM potassium bicarbonate for 144 hours (37°C, pH 7.0).
<sup>b</sup> Fermentations (n ≥ 3) with 0.1 mM IPTG added to induce the expression of candidate genes.
<sup>c</sup> Yield was calculated as g malate produced per g glucose consumed.
<sup>d</sup> All data at 144 hours were shown here and SD were included. Abbreviations: Mal, malate; Lac, lactate; Pyr, pyruvate; Ace, acetate; For, formate.

Plasmid-based systems have several disadvantages in large-scale microbial production, such as genetic instability, performance inconsistency, metabolic burden and the requirement of antibiotics and inducers (Jarboe et al., 2010; Ow et al., 2006). We integrated *citZ* from both *Bacillus* strains into the *frdBC* locus on the chromosome since the *frdBC* operon was already deleted in the malate producing strain XZ658 and the upstream promoter is active under anaerobic fermentation conditions (Ruch et al., 1979). GltA R163L was not used for chromosomal integration due to potential homologous recombination to restore the mutation and cause undefined genetic changes. Integration of each *Bacillus* citZ gene increased cell growth and malate production metrics, with *B. subtilis* citZ slightly outperforming *B. licheniformis* citZ in terms of malate titer and yield (data not shown). The integration of *B. subtilis citZ* yielded the strain JB02 that produced 33 g liter^-1^ with a product yield of 0.9 g g^-1^ glucose by a simple batch fermentation. Compared to the precursor strain XZ658, this genetic modification increased the titer, yield, and productivity by 50%, 20%, and 50%, respectively (Fig. 2). In addition, the replacement of native *E. coli gltA* with *B. licheniformis citZ* in XZ658 (JB03) also showed a similar positive effect, suggesting that type I citrate synthase can functionally replace native type II enzyme in XZ658 without noticeable growth defect under fermentation conditions (data not shown).

**Figure 2.**
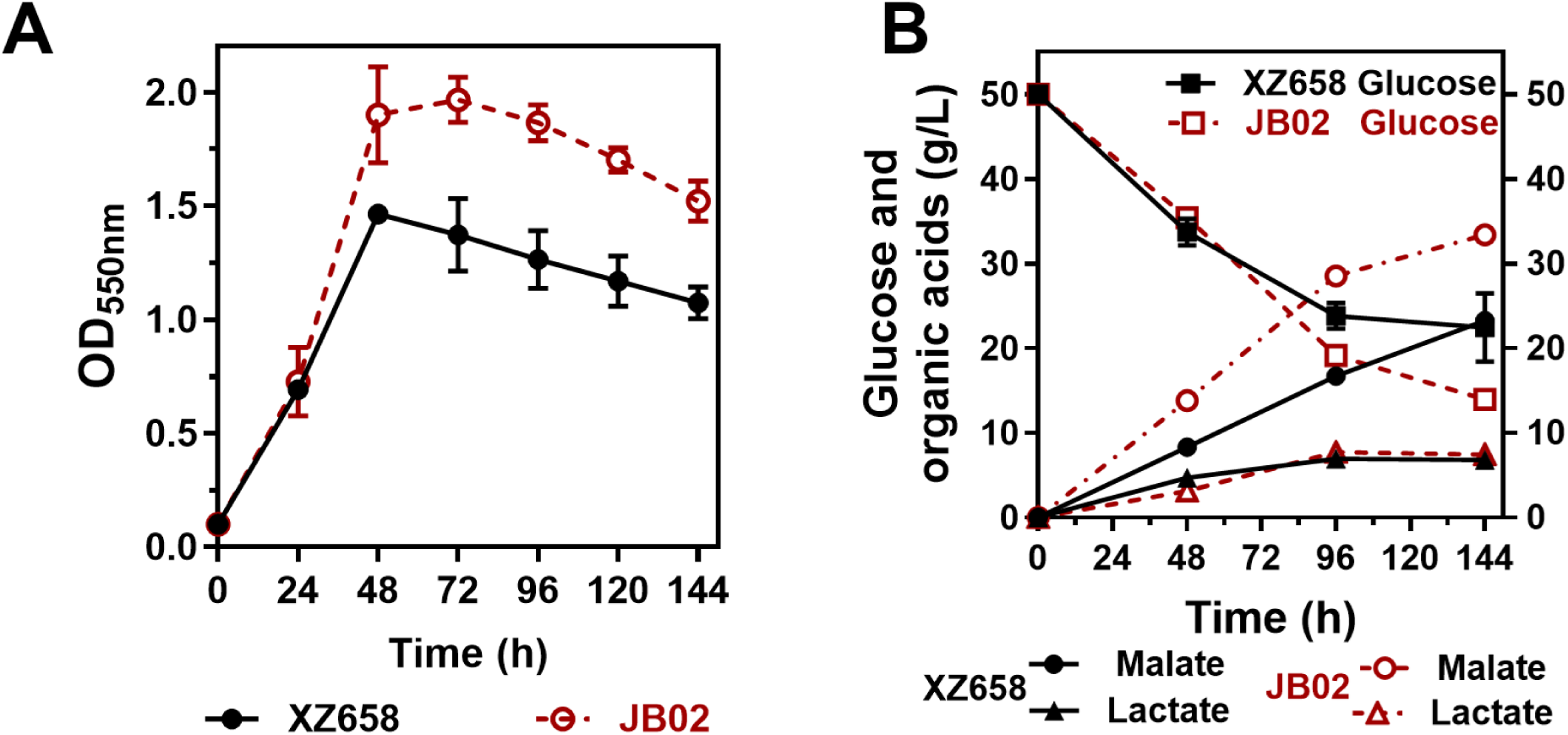
Fermentation profile of XZ658 and JB02 using mineral salts media containing 50 g liter^-1^ glucose. A) Growth curve of XZ658 and JB02 measured at OD_550nm_. B) Glucose consumption, malate production, and lactate byproduct production of XZ658 compared to JB02. Symbols: black solid lines and red dotted lines represent XZ658 and JB02 fermentation profiles, respectively.

### 3.2. Metabolite analysis reveals NADH/NAD^+^ imbalance as a potential constraint for malate overproduction

Allosteric inhibition of citrate synthase by NADH in *E. coli* may reduce intracellular levels of α-ketoglutarate, an important precursor for amino acid biosynthesis, thereby negatively affecting cell growth, particularly under oxygen-limited fermentation conditions with elevated NADH/NAD⁺ ratios (Underwood et al., 2002). Comparison of intracellular metabolites between JB02 and XZ658 revealed increased levels of metabolites in the oxidative branch of the TCA cycle in JB02, including citrate (2-fold), aconitate (3.4-fold), isocitrate (3-fold), and α-ketoglutarate (2.2-fold), as shown in Fig. 3A and Fig. 3B. Metabolites relevant to malate biosynthesis, such as PEP, pyruvate, and malate, were also increased by several fold, consistent with the observed improvement in malate yield. These changes in metabolite levels favor fermentative malate production.

**Figure 3.**
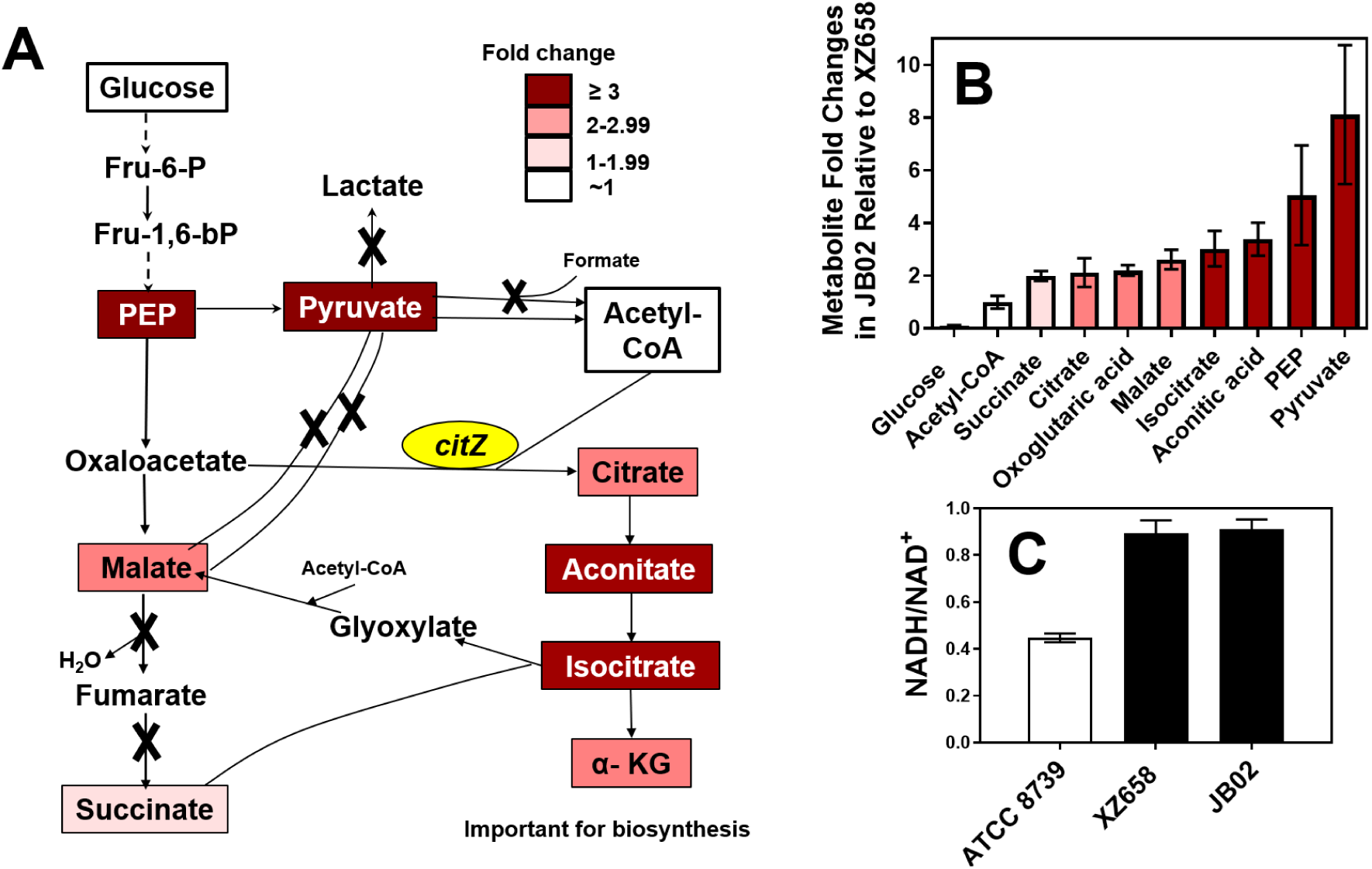
Releasing allosteric regulation of citrate synthase alters intracellular metabolite levels. A) Schematic of the redox-balanced malate-producing pathway using the reductive TCA branch; deleted genes are indicated by “X.” Introduction of *citZ* alters the relative abundance of pathway metabolites. B) Relative fold changes of intracellular metabolites in JB02 versus XZ658 measured by GC–MS; mean values normalized by protein concentration compared to XZ658 means. Error bars represent SEM (n ≥ 3). C) Intracellular NADH/NAD⁺ ratios in malate-producing strains compared to wild-type *E. coli*. Abbreviations: Fru-6-P, fructose-6-phosphate; Fru-1,6-bP, fructose-1,6-bisphosphate; PEP, phosphoenolpyruvate; α-KG, α-ketoglutarate; acetyl-CoA, acetyl-Coenzyme A.

The observed allosteric regulation of citrate synthase under high NADH conditions suggests that the NADH/NAD⁺ ratio exceeds the optimal level. Indeed, intracellular NADH/NAD⁺ measurements (Fig. 3C) showed that both malate-producing strains, XZ658 and JB02, exhibited ∼1.25-fold higher ratios compared to wild-type (WT) *E. coli* ATCC 8739 (the parental strain of the malate-producing strains) under fermentative conditions. Because NADH inhibits several key enzymes involved in biosynthesis and malate production, including the pyruvate dehydrogenase complex, isocitrate dehydrogenase, α-ketoglutarate dehydrogenase complex, and malate dehydrogenase (Sanwal, 1969), these results indicate that NADH/NAD⁺ imbalance remains a potential constraint on malate overproduction in JB02. Reducing the NADH/NAD⁺ ratio may therefore enhance biosynthetic activity and improve malate fermentation.

### 3.3. Reducing NADH/NAD^+^ ratio to enhance malate production

Under oxygen-limited fermentation conditions, a high NADH/NAD⁺ ratio indicates low efficiency of the downstream fermentation pathway, specifically the malate dehydrogenase (MDH)-catalyzed reduction, since all other fermentation pathways have been deleted in JB02 (Fig. 1A). MDH catalyzes the reduction of oxaloacetate to malate using NADH as a cofactor, thereby regenerating NAD⁺. To increase NAD⁺ regeneration and reduce the NADH/NAD⁺ ratio in JB02, we overexpressed selected malate dehydrogenase genes (MDHs) from different microorganisms. In addition to the native MDH, enzymes from *Synechocystis* sp. PCC 6803 and *Corynebacterium glutamicum* were chosen based on their reported higher activities toward oxaloacetate reduction (Ahn et al., 2020; Takeya et al., 2018).

First, the native *mdh* gene was deleted in the XZ658 background, and the resulting strain was transformed with plasmids harboring the MDH genes to validate their functional expression (data not shown). After confirming functional expression by restored growth and malate production, the MDH plasmids were introduced into JB02 to evaluate their effects. None of the tested MDHs improved fermentation performance compared to the empty vector control in JB02 (Fig. 4). These results suggest that simply increasing MDH activity is not sufficient to resolve redox imbalance in malate-producing strains.

**Figure 4.**
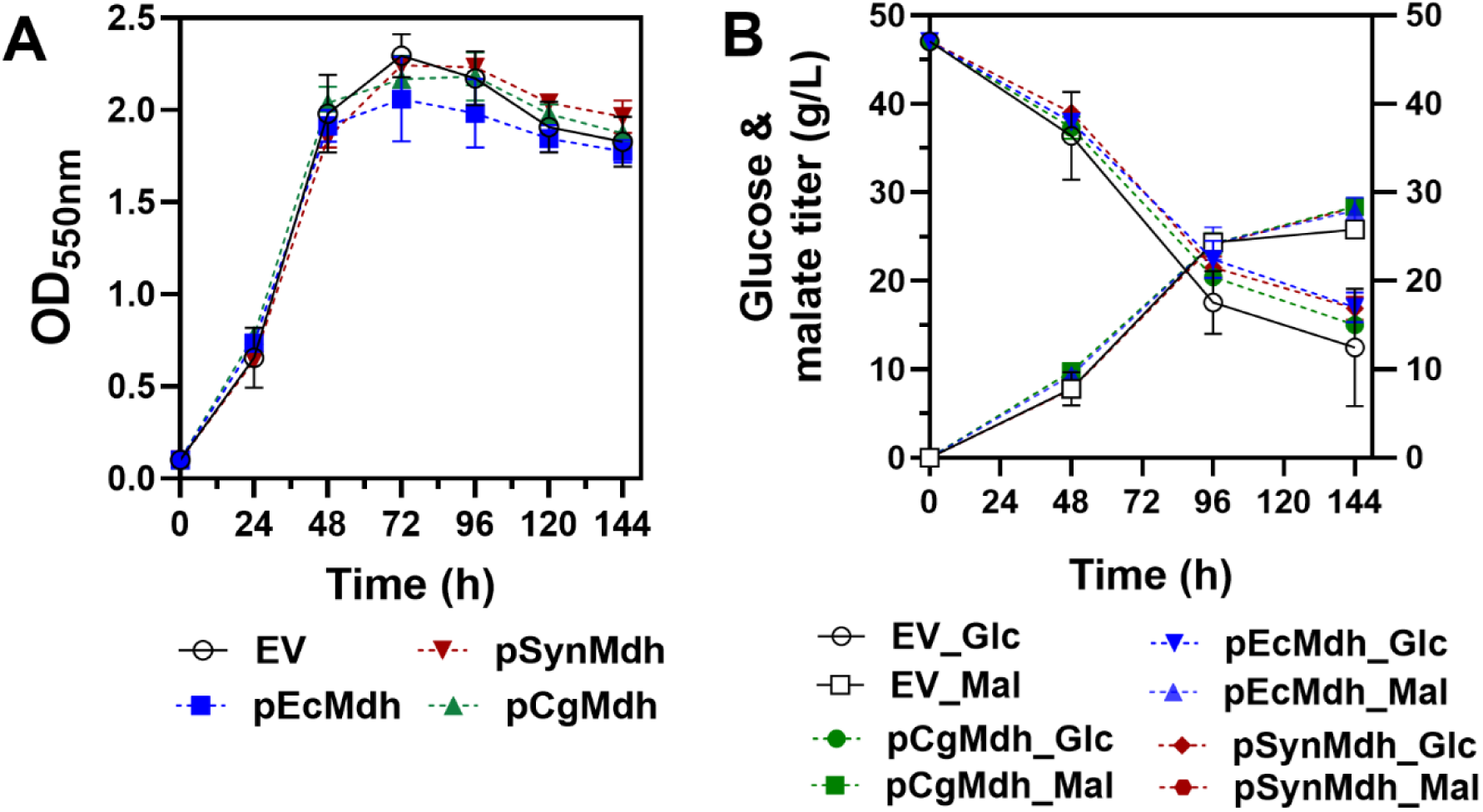
Fermentation profiles of JB02 expressing malate dehydrogenase isozyme genes. A) Growth curves of JB02 expressing different *mdh* genes. B) Effects of plasmid-based *mdh* expression on glucose consumption and malate production in JB02. Abbreviations: EV, empty vector; EcMdh, *E. coli* malate dehydrogenase; CgMdh, *C. glutamicum* malate dehydrogenase; SynMdh, *Synechocystis* sp. PCC 6803 malate dehydrogenase; Mal, malate; Glc, glucose; Lac, lactate.

We sought to address the high NADH/NAD⁺ ratio by introducing an alternative electron acceptor into the medium. Many bacteria, including *E. coli*, can use nitrate as an alternative electron acceptor under anaerobic conditions (Stewart, 1988). The reduction of nitrate to nitrite by nitrate reductase enables NADH oxidation (Fig. 5A). Therefore, we evaluated the effect of nitrate supplementation on cell growth and malate production in JB02. Nitrate salts are inexpensive, easily incorporated into culture media, and *E. coli* possesses the native genes required for anaerobic nitrate respiration (Fig. 5A) (Stewart, 1988). We tested the effects of different nitrate concentrations (5, 10, 15, and 20 mM) on fermentative performance. Supplementation with 5 mM potassium nitrate increased maximal cell growth and malate titer by 20% and 24%, respectively. Similarly, 10, 15, and 20 mM potassium nitrate increased maximal cell growth by 32.5%, 66.7%, and 81%, and malate titer by 24%, 30%, and 30%, respectively (Fig. 5B and 5C). Notably, lactate, the main byproduct, decreased by approximately 50% upon nitrate supplementation (Fig. 5D), resulting in an increased yield (∼1.1 g g⁻¹ glucose with 15 mM KNO₃).

**Figure 5.**
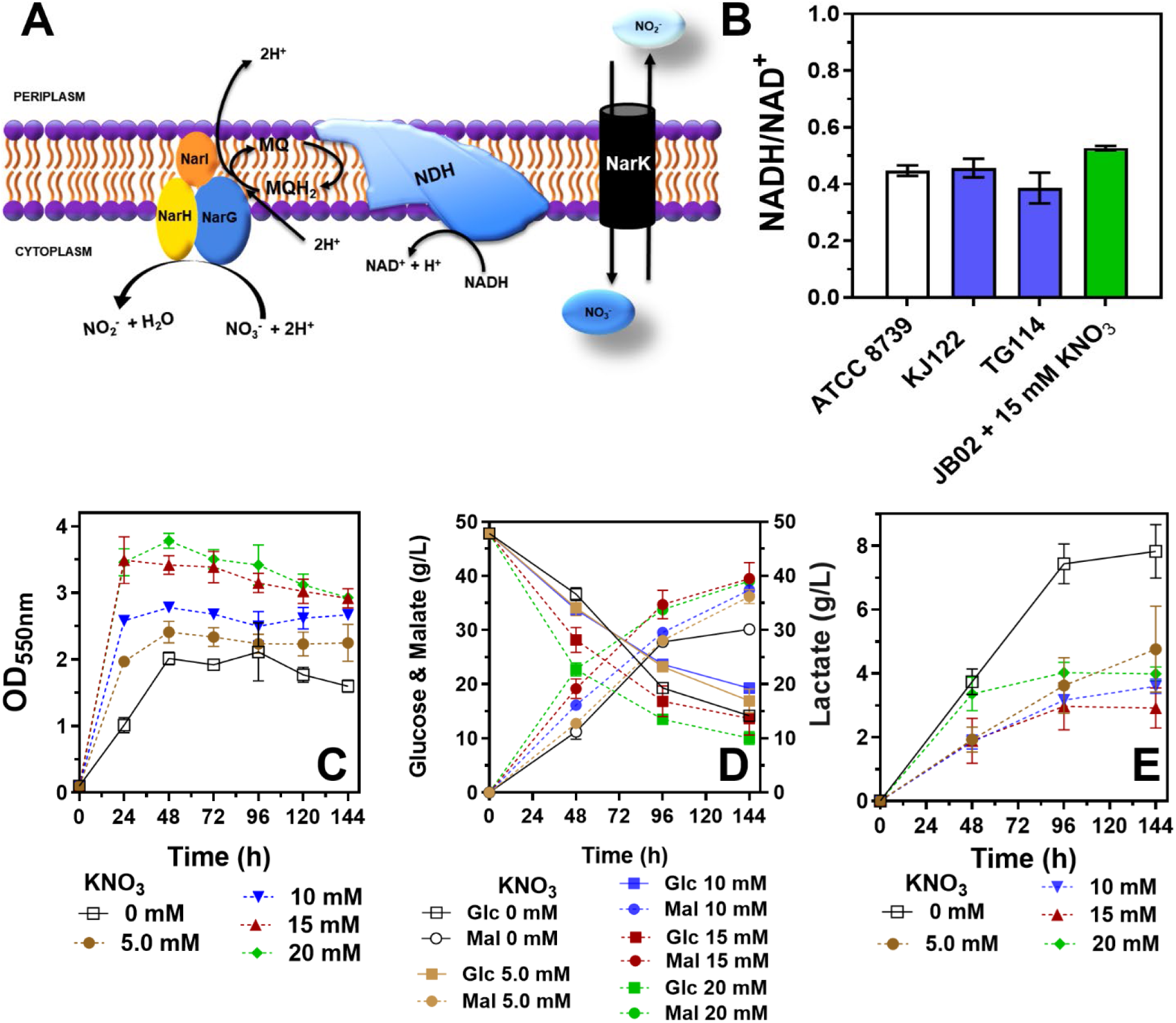
Nitrate supplementation improves growth and malate production. A) Simplified schematic of *E. coli* nitrate reduction, coupled to NADH oxidation. B) Intracellular NADH/NAD⁺ ratios of JB02 with 15 mM nitrate, compared to wild-type ATCC 8739 and KJ122. Data were obtained from cultures in the late exponential growth phase under fermentation conditions. C) Effect of nitrate concentration on cell growth. D) Effect of nitrate on glucose consumption and malate production. E) Lactate production decreases with nitrate supplementation. Cells were fermented in NBS medium with 5% (w/v) glucose, with or without nitrate.

Although biomass accumulation was higher with 20 mM nitrate (81%) than with 15 mM nitrate (66.7%), the final malate titer was similar in both cases (∼40 g L⁻¹). Based on these results, 15 mM nitrate was selected for subsequent experiments.

Next, we evaluated whether nitrate supplementation effectively reduces the NADH/NAD⁺ ratio in the fermentation medium. Indeed, with 15 mM nitrate supplementation, the NADH/NAD⁺ ratio decreased by 42% to 0.53 compared to the condition without nitrate (Fig. 3C and Fig. 5E). As shown in Fig. 5E, the NADH/NAD⁺ ratio of JB02 with 15 mM nitrate is comparable to that of the wild-type strain and other engineered *E. coli* strains optimized for fermentative succinate (KJ122) and D-lactate production (TG114) (Grabar et al., 2006; Jantama et al., 2008).

The reduced NADH/NAD⁺ ratio is consistent with improved biomass accumulation and malate production, highlighting the importance of redox balance in microbial fermentation and bioproduction. In addition, the elevated NADH/NAD⁺ ratio appears to promote lactate accumulation in XZ658, likely as a route for NADH oxidation. However, the exact mechanism remains unclear, particularly since the lactate dehydrogenase gene is deleted in JB02. A more balanced redox state likely redistributes metabolic flux, leading to higher malate yield and titer. Overall, nitrate supplementation at appropriate concentrations represents an effective strategy to lower the NADH/NAD⁺ ratio and achieve improved redox balance.

### 3.4. Enhancing malate export capacity improves malate production under nitrate-supplemented conditions

We previously identified three transporters (DcuA, CitT, and TtdT) as major components of the malate export system in *E. coli*, with DcuA serving as the primary transporter (Kurgan et al., 2019). However, overexpression of *dcuA*, either from a plasmid or from the chromosome, did not increase malate titer in XZ658(Kurgan et al., 2019). Here, we evaluated whether malate export limits overproduction in JB02. Overexpression of *dcuA* in JB02 did not improve malate production in the absence of nitrate (Fig. 6A and 6B), suggesting that allosteric inhibition and redox imbalance may be more dominant constraints. Interestingly, in the presence of 15 mM KNO₃, cells expressing *dcuA* induced with 10 µM IPTG showed improved growth compared to the empty vector control, resulting in a 9% increase in malate titer (from 34 g L⁻¹ to 37 g L⁻¹) (Fig. 6C and 6D). Increasing IPTG to 100 µM produced a similar effect, with an 18% increase in titer (from 34 g L⁻¹ to 40 g L⁻¹). Notably, lactate production decreased by 31% and 14% upon *dcuA* overexpression with 10 µM and 100 µM IPTG, respectively. Enhanced malate export may redistribute metabolic flux, leading to reduced lactate formation. In addition, nitrate supplementation may improve bioenergetic conditions, including the proton motive force, thereby facilitating malate export.

**Figure 6.**
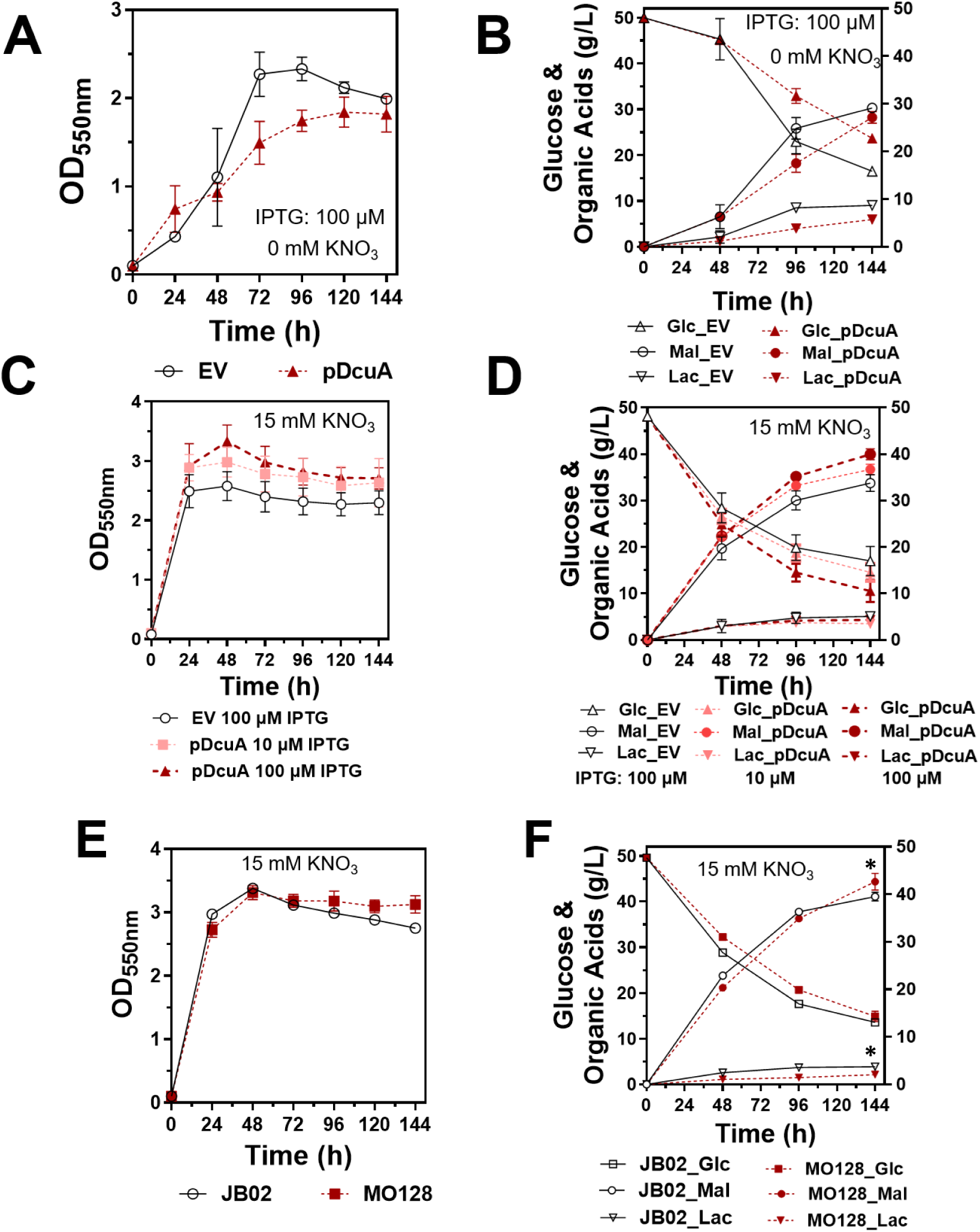
Effect of *dcuA* overexpression on growth and L-malate production. Without nitrate, A) cell growth and B) glucose consumption as well as organic acids of JB02 harboring pDcuA or empty vector (EV) were measured. 100 µM IPTG was used to induce *dcuA* expression. In the presence of 15 mM KNO_3_, the effects of pDcuA on C) cell growth and D) glucose consumption, malate production as well as lactate formation were measured. 10 µM and 100 µM IPTG were used to induce *dcuA* expression while 100 µM IPTG was used for EV control. The impact of extra chromosomal *dcuA* expression in MO128 (*ldhA*::*dcuA* and *adhE*::*dcuA*) on E) cell growth as well as F) glucose consumption, malate and lactate production were measured. Cells were fermented in NBS medium with 5% (w/v) glucose with 15 mM KNO_3_. Abbreviations: Mal, malate; Glc, glucose; Lac, lactate. Asterisks indicate statistical significance by Student’s t-test (p < 0.05); however, the small sample size (n = 3) warrants cautious interpretation.

Next, we integrated an additional copy of *dcuA* into the chromosome of JB02. The *ldhA* and *adhE* loci were selected because the corresponding genes are deleted and their promoters are active under fermentation conditions. The resulting strains, MO122 (*ldhA::dcuA*) and MO124 (*adhE::dcuA*), did not show improved malate titers (Fig. S1). However, consistent with the phenotype observed for plasmid-based expression, both MO122 and MO124 exhibited reduced D-lactate titers by 52% and 63%, respectively (Fig. S1E).

Since malate titer was highest when *dcuA* expression was induced with 100 µM IPTG from a high-copy plasmid, we reasoned that a single chromosomal integration may be insufficient to achieve the required expression level. Consequently, *dcuA* was integrated into both the *ldhA* and *adhE* loci to generate strain MO128. Although the growth profile of MO128 was comparable to that of JB02 at the beginning of fermentation, MO128 maintained slightly higher biomass toward the end of fermentation (Fig. 6E). Accordingly, MO128 showed a modest improvement in malate titer, reaching 44.3 g L⁻¹. In addition, D-lactate production decreased by 45% in MO128, corresponding to an increase in malate yield from 1.1 g g⁻¹ in JB02 (with nitrate supplementation) to 1.2 g g⁻¹ in MO128, representing a 9% improvement.

## 4. Discussion

Fermentative L-malate production via the reductive branch of the TCA cycle has the potential to assimilate inorganic carbon and achieve higher yields (1.49 g malate g⁻¹ glucose; 2.0 mol mol⁻¹) than conventional fermentation products such as ethanol (0.51 g g⁻¹) and lactate (1.0 g g⁻¹). However, as is common for non-native fermentation products, multiple native metabolic constraints limit malate overproduction. In this study, we identified allosteric regulation of citrate synthase, an imbalanced NADH/NAD⁺ ratio, and suboptimal export as key factors limiting malate production and cell growth in *E. coli*. Addressing these constraints approximately doubled the L-malate titer in the final strain MO128 under simple batch fermentation conditions. The process achieved a high mass yield of 1.2 g malate g⁻¹ glucose, highlighting the carbon-fixation capacity of the reductive TCA pathway for fermentative malate production.

Allosteric regulation of key enzymes is one of the primary mechanisms controlling complex cellular metabolic networks. Identification and deregulation of these biochemical controls have been shown to improve microbial production, particularly for non-native fermentation products such as amino acids and aromatic compounds (Frost and Draths, 1995; Wendisch, 2014). Relief of allosteric regulation of citrate synthase in the malate-producing strain not only enhanced cell growth but also significantly altered intracellular metabolite levels (Fig. 3A and 3B). These metabolic changes increased specific malate production on a per-cell basis.

The increased malate production is unlikely to result from an enhanced glyoxylate shunt, as succinate production in the engineered strains (JB01, JB02, and JB03) remained negligible (<1 g L⁻¹; data not shown) compared to the elevated malate titers (∼10–13 g L⁻¹; Fig. 2). Relief of citrate synthase regulation increased intracellular levels of citrate and α-ketoglutarate, which are important for amino acid biosynthesis and other physiological processes (Fig. 3A and 3B). A similar observation was reported in ethanologenic *E. coli* KO11 (Underwood et al., 2002), where relief of citrate synthase allosteric regulation improved fermentative growth and ethanol production when xylose was used as the sole carbon source.

Our study showed that the NADH/NAD⁺ ratio is critical for efficient fermentation. Wild-type and engineered *E. coli* strains with efficient fermentative performance exhibited relatively lower NADH/NAD⁺ ratios compared to XZ658 or JB02, indicating more efficient fermentation pathways and/or product export. We attempted to improve the malate fermentation pathway through overexpression of MDH genes and adaptive laboratory evolution; however, both approaches were unsuccessful (Fig. 4 and data not shown).

As an alternative strategy, we used nitrate as an external electron acceptor to reduce the NADH/NAD⁺ ratio. *E. coli* possesses the necessary enzymes and transport systems for nitrate metabolism. Nitrate reduction oxidizes NADH and generates a proton gradient that can be utilized for ATP synthesis via ATP synthase or proton-dependent product export (Stewart, 1988). In addition to redox balancing and ATP generation, nitrate respiration has been reported to increase flux through the pentose phosphate pathway, thereby enhancing NADPH supply for anabolic processes (Toya et al., 2012). Collectively, these metabolic effects likely contribute to the observed improvement in fermentative malate production. We are not aware of any previous study where nitrate is used to optimize NADH/NAD^+^ ratio for enhanced malate microbial production. This *E. coli*-based fermentative process, requiring only minimal media and low nitrate concentrations, offers a simple and potentially scalable route for biotechnological L-malate production.

D-lactate is the main byproduct in the malate-producing strain XZ658 (Zhang et al., 2011). Lactate is likely derived from pyruvate via an as-yet-unidentified metabolic route. It is plausible that lactate formation supports cell growth by oxidizing excess reducing equivalents generated during glycolysis, thereby contributing to redox balance. Nitrate supplementation enhances NADH oxidation, which relieves the requirement for NADH reoxidation through D-lactate formation. As a result, lactate production decreased by approximately 50% in JB02 with nitrate supplementation (Fig. 5E). Although the exact mechanism of lactate accumulation remains unclear, this byproduct formation was unexpectedly but effectively reduced by nitrate supplementation.

Although overexpression of a native C4-dicarboxylate transporter encoded by the *C4T318* gene led to more than a two-fold increase in malate production rate in *Aspergillus oryzae* NRRL 3488 (Brown et al., 2013), and overexpression of the *Schizosaccharomyces pombe* malate transporter gene *SpMAE1* promoted malic acid production in engineered *Saccharomyces cerevisiae* (Zelle et al., 2008), overexpression of *dcuA* in JB02 did not improve malate production in the absence of nitrate (Fig. 6A and 6B). This result is consistent with our previous *dcuA* overexpression study in XZ658. It suggests that either malate export is not a primary constraint or that other limiting factors dominate over export capacity.

Interestingly, in the presence of nitrate supplementation, overexpression of *dcuA* led to a modest improvement in malate production, particularly in yield (Fig. 6). This suggests that malate production may require substantial energy input, which cannot be resolved by transporter overexpression alone. Instead, optimizing cellular bioenergetics appears to be a key requirement for efficient fermentative production.

There is still room for improvement in L-malate production using *E. coli*, particularly in terms of titer, compared with fungal production systems producers (Wu et al., 2025; Xu et al., 2024). In addition to cellular bioenergetics, remaining bottlenecks in MO128 may include suboptimal performance of key carbon-fixation enzymes involved in malate production. Phosphoenolpyruvate carboxylation is an important step in malate biosynthesis in *E. coli* (Fig. 1A). Optimizing this carbon-fixation step may therefore further enhance both cell growth and malate production.

