## Supplementary material for "Relief of allosteric inhibition, redox imbalance, and transport limitations enables high-yield L-malate production in *Escherichia coli*": Table S1 and Fig. S1

### Supplemental Information

**Table S1**. Primers used in this study.

| | Sequences | Reference |
|---|---|---|
| *pfkA* T125A cloning | CGACGGTACCCAAGAAGACTTCCGGCAACAG | This study |
| | ATGGTCTAGAGTTGAGGGATTAAAAAGGCGG | |
| | CCGTGCATCGGTCTGCCGGGCTCTATCGACAACGACATC | |
| | CCTTTGATGTCGTTGTCGATAGAGCCCGGCAGACCGATG | |
| *Lpd* E354K cloning | CGTACGAATTCGAAAGACGACGGGTATGACCG | This study |
| | GTGTTCTAGACGACTGGAAAGGTAAATTACAGACG | |
| | ATCGCCTATACCAAACCAGAAGTTG | |
| | CAACTTCTGGTTTGGTATAGGCGAT | |
| *gltA* R163L cloning | CGTACGAATTCGTTCCGGCAGTCTTACGC | This study |
| | GTGTTCTAGAGTTCAGCCATATAAAAAGAACCCGC | |
| | GCCGCGTTCCTCCTGCTGTCGA | |
| | TCGACAGCAGGAGGAACGCGGC | |
| *citZ*$_{\mathit{B.\ subtilis}}$ cloning | AGAAGTGGGATCCTGGGGGAGAGAAATACTTGC | This study |
| | TTAATGGGTCGACTCGGGTTGTTTGGTACGTTT | |
| *citZ*$_{\mathit{B.\ licheniformis}}$ cloning | GAAGTGGGATCCCAACATGCGTTGTTTTTGCTCC | This study |
| | TTAATGGGTCGACACATTTGTTTCCCCCGTTTG | |
| *mdh*$_{\mathit{E.\ coli}}$ cloning | ACACAGGAAACAGACCATGGAATTCCGGCAGCGGAGCAACATATC | This study |
| | ACAACTGTCGCAAAAATAATGTTCAAAAACAAAAAACCGGAG | |
| | CTCCGGTTTTTTGTTTTTGAACATTATTTTTGCGACAGTTGT | |
| | GATATGTTGCTCCGCTGCCGGAATTCCATGGTCTGTTTCCTGTGT | |
| *Mdh*$_{\mathit{Synechocystis}}$ cloning | CACAGGAAACAGACCATGGAATTCCCGCCTGACATTGTTGAATTTCG | This study |
| | TGTCGCAAAAATAATGTTCAACTATTGACCGGCCACCCCTTC | |

| | | |
|---|---|---|
| | GAAGGGGTGGCCGGTCAATAGTTGAACATTATTTT<br>TGCGACA<br>CGAAATTCAACAATGTCAGGCGGAATTCCATGGT<br>CTGTTTCCTGTG | |
| *Mdh*$_{C.\ glutamicum}$ cloning | ATTTCACCTGCTTGAGCTCAGCGGGATAGCATGGG<br>TTC<br>ATTTCACCTGCTTGACTATATTAGAGCAAGTCGCG<br>CAC | This study |
| *dcuA*$_{E.\ coli}$ cloning | ATTTCACCTGCTTGAGCTCAAAGAAGGCACGTCAG<br>AT<br>ATTTCACCTGCTTGACTATAGGTCACTATACAACG<br>GAC | This study |
| *frdBC::cat-sacB integration* | AGCGGATGCAGCCGATAAGGCGGAAGCAGCCAAT<br>AAGAAGGAGAAGGCGATCGAGTGTGACGGAAGAT<br>CA<br>ATACCGGTTCGTCAGAACGCTTTGGATTTGGATTG<br>ATCATCTCAGGCTCCTTAGCCATTTGCCTGCTTTT<br>TATCTACCGTACGCCGGAAC (used for sequencing)<br>AGCAAATGTGGAGCAAGAGG (used for sequencing) | This Study |
| *frdBC::citZ*$_{B.\ licheniformis}$ *and frdBC::citZ*$_{B.\ subtilis}$ *integrations* | AGCGGATGCAGCCGATAAGGCGGAAGCAGCCAAT<br>AAGAAGGAGAAGGCGAATGACAGCGACACGCGGT<br>C<br>ATACCGGTTCGTCAGAACGCTTTGGATTTGGATTG<br>ATCATCTCAGGCTCCTTAGGCTCTTTCTTCAATCGG<br>AACGAA | This study |
| *gltA::cat-sacB* | AAATTTAAGTTCCGGCAGTCTTACGCAATAAGGCG<br>CTAAGGAGACCTTAATCGAGTGTGACGGAAGATCA<br>CCCGCCATATGAACGGCGGGTTAAAATATTTACAA<br>CTTAGCAATCAACCATTAGCCATTTGCCTGCTTTT | This study |
| *gltA::citZ*$_{B.\ licheniformis}$ *and gltA::citZ*$_{B.\ subtilis}$ *integrations* | AAATTTAAGTTCCGGCAGTCTTACGCAATAAGGCG<br>CTAAGGAGACCTTAAATGACAGCGACACGCGGTC<br>CCGCTCTATTAAAGGCGGGTCCGGAAAGTAAACGG<br>CTTAGCAATCAATCATTAGGCTCTTTCTTCAATCGG<br>AACGAA | This study |

| | | |
|---|---|---|
| *ldhA::cat-sacB integration* | AAATATTTTTAGTAGCTTAAATGTGATTCAACATCACTGGAGAAAGTCTTATCCTGGTGTCCCTGTTGATACCG<br>ATTGGGGATTATCTGAATCAGCTCCCCTGGGTTGCAGGGGAGCGGCAAGATGGCCGATTCATTAATGCAGCTGG | This study |
| *ldhA::dcuA integration* | ACCAGCGGCTGGGATGTGAAAG<br>CATCTGACGTGCCTTCTTTAAGACTTTCTCCAGTGATGTTG<br>GTCCGTTGTATAGTGACCTTCTTGCCGCTCCCCTGCAAC<br>AGCTGCGGGTTAGCGCACATC<br>CAACATCACTGGAGAAAGTCTTAAAGAAGGCACGTCAGATG<br>GTTGCAGGGGAGCGGCAAGAAGGTCACTATACAACGGAC | This study |
| *adhE::cat-sacB integration* | ATTCGAGCAGATGATTTACTAAAAAAGTTTAACATTATCAGGAGAGCATTATCCTGGTGTCCCTGTTGATACCG<br>ATCGGCATTGCCCAGAAGGGGCCGTTTATGTTGCCAGACAGCGCTACTGATGGCCGATTCATTAATGCAGCTGG | This study |
| *adhE::dcuA integration* | AAAGGTCTGAATCACGGTTAG<br>CATCTGACGTGCCTTCTTTAATGCTCTCCTGATAATGTTAAAC<br>GTCCGTTGTATAGTGACCTAATCAGTAGCGCTGTCTG<br>AGTCATCCTTCAGGTAACG<br>GTTTAACATTATCAGGAGAGCATTAAAGAAGGCACGTCAGATG<br>CAGACAGCGCTACTGATTAGGTCACTATACAACGGAC | This study |

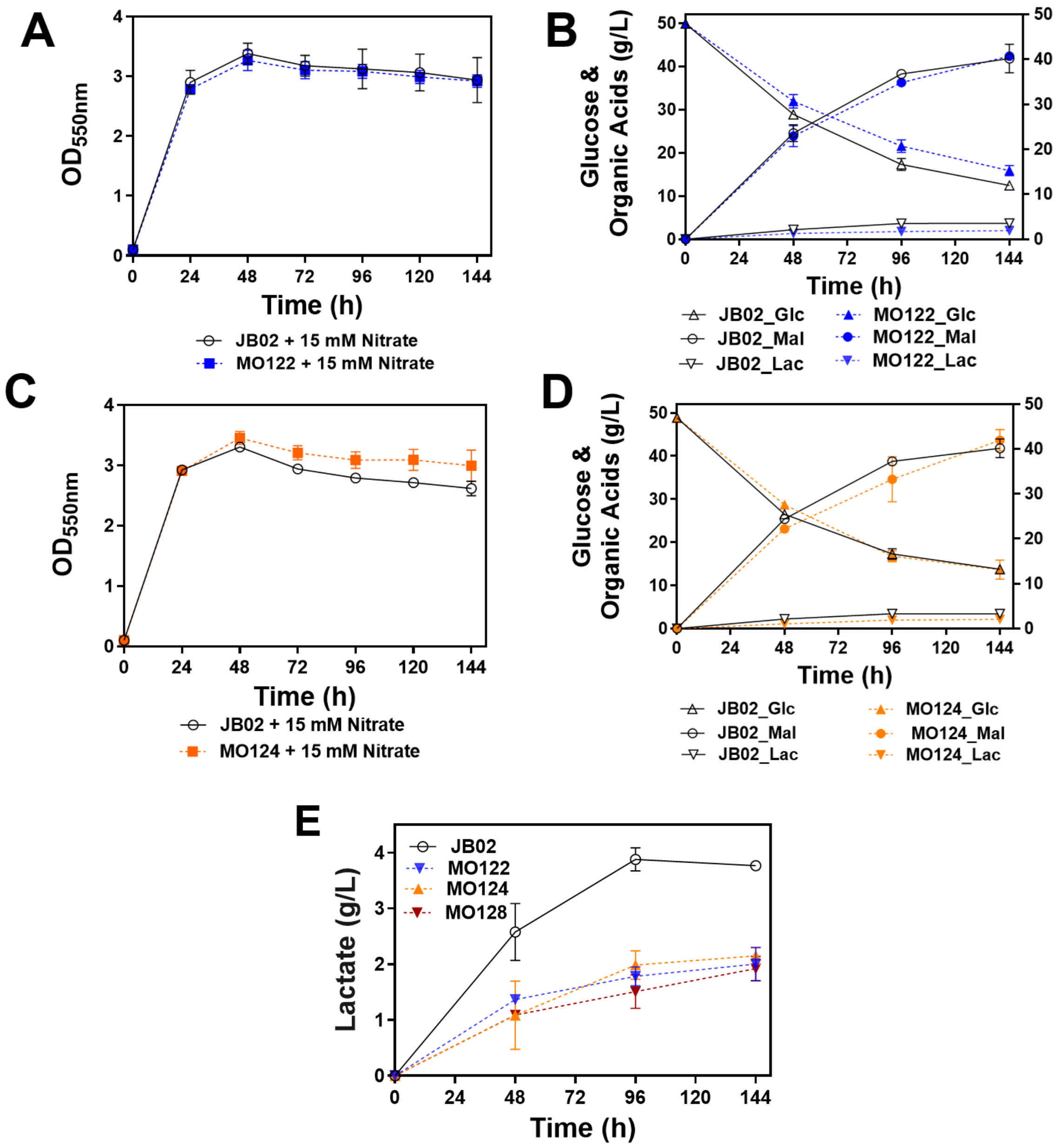


**Figure S1**. Single *dcuA* integration into either *ldhA* (MO122) or *adhE* (MO124) did not improve malate production. A) Growth curve of MO122 and JB02. B) Effect of *dcuA* integration into *ldhA* locus on glucose consumption, malate, and byproduct lactate production. C) Growth curve of MO124 and JB02. D) Effect of *dcuA* integration into *adhE* locus on glucose consumption, malate, and byproduct lactate production. E) Effect of *dcuA* chromosomal integration on lactate production for MO122, MO124 and MO128 (*dcuA* integrated in both *ldhA* and *adhE* loci). Cells were fermented in NBS 5% glucose (w/v) medium with 15 mM $KNO_3$ supplementation. Abbreviations: Mal, malate; Glc, glucose; Lac, lactate.
